# Five Rotational Registers Define Modular Portal-Capsid Assembly in Cyanophage P-SSP7

**DOI:** 10.64898/2026.08.19.745834

**Authors:** Amar D. Parvate, Pavlo Bohutskyi, Natalie Sadler, Margaret S. Cheung, James E. Evans

## Abstract

The virus P-SSP7 infects the cyanobacterium *Prochlorococcus marinus*, one of the most abundant photosynthetic microbes in the ocean, making this pairing a useful model for studying host-virus interactions. Infection proceeds through a portal-tail complex that attaches to the viral protein shell at a single specialized vertex, where the two structures have mismatched symmetries. Previous studies described this region only at coarse resolution which continues to limit understanding of how the virus assembles and injects its genome. Using cryo-electron microscopy of high-quality samples, we determined the first near-atomic-resolution structure of the complete P-SSP7 virion and its portal-tail complex. We captured 3D classes where the shell and portal-tail complex connect in five distinct arrangements. Fitting and modeling 2 of these states helped resolve and establish modular attachment in five rotational registers as the basis for capsid-portal symmetry mismatch in P-SSP7 assembly.

## Introduction

Picocyanobacteria such as *Prochlorococcus marinus* are key drivers of marine photosynthesis and primary production^1,2^. Cyanophages prey on these hosts; lysis releases fixed carbon and nitrogen into seawater and regulates cyanobacterial populations, metabolic turnover, and nutrient cycling^3,4^. Beyond these ecological roles, cyanophages shape host metabolism and evolution through horizontal exchange of auxiliary metabolic genes (AMGs) spanning photosynthesis, central carbon metabolism, and nutrient stress responses^5–10^. These functions ultimately depend on virion architecture, particularly the portal–tail assembly that mediates host attachment and genome delivery, motivating high-resolution structural analysis of intact cyanophages.

Cyanophage P-SSP7 is a podovirus that infects *Prochlorococcus marinus* and serves as a model system for host–pathogen interactions in marine picocyanobacteria. Its genome, transcriptome, and proteome, and the corresponding host profiles during infection, have been characterized^8,11^. Similarities to the genome organization of T7 suggest that P-SSP7 uses replication and packaging strategies related to T7-like phages^11^. Cryo-EM and cryo-ET established the overall P-SSP7 architecture, including a non-enveloped, ∼655 Å diameter capsid arranged in a T = 7 icosahedral lattice^3,5^.

Of the twelve five-fold vertices on the icosahedral capsid, one is unique and contains the portal complex, which functions as an entry/exit channel for the genome. In prevailing models, scaffold proteins guide procapsid formation and a 12-fold portal assembles at the unique 5-fold vertex; a protein motor then packages dsDNA through this vertex^12^. Following packaging, the motor dissociates and the tail nozzle attaches to the portal vertex to form the mature phage^13^. The interaction of a 12-fold portal with a 5-fold capsid vertex creates a symmetry mismatch common across icosahedral viruses, including mammalian viruses, phages, and archaeal viruses. In tailed phages, this mismatch is particularly notable because capsid and portal complexes differ not only in structure but also in biological function. During infection, tailed phages adsorb to the host surface and inject the stably packaged genome into the cell to initiate replication^13^; these steps require large-scale macromolecular rearrangements. Models propose that the symmetry mismatch at the capsid–portal vertex provides flexibility needed for genome retention during assembly and genome release during infection^14^. Since the initial reports over 25 years ago^15^, researchers have described scaffold flexibility (HK97)^16^, connector rotation (phi29)^15,17^, non-integral/uncorrelated arrangements (T7)^18^, continuous morphing (T4)^19^, breaking 12-fold symmetry of the portal complex (P-SCSP1u)^20^ and two equivalent tail conformations (SF6 and P22)^21,22^ as the reported solutions for the 12:5 symmetry mismatch.

Multiple studies have analyzed P-SSP7 by Cryo-EM and Cryo-ET as purified particles and during infection^3,5^. However, the Protein Data Bank contains only a 4.6 Å capsid map and a nanometer-resolution portal–tail map. The current gap in our knowledge is atomic-resolution structural information across the virion, which is needed to dissect assembly, host interaction, and genome delivery. The 12:5 portal–capsid mismatch, together with a milder 12:6 portal–nozzle mismatch, limits symmetry averaging and demands substantially larger datasets for high-resolution cryo-EM single-particle reconstructions of the portal–tail complex.

Here, we leveraged our optimized cyanophage sample preparation^23^ to collect a large cryo-EM dataset (32,000 movies, >1 million particles) to address these challenges. From this dataset we obtained a 3.02 Å reconstruction of the entire P-SSP7 phage. 3D classification identified 5 distinct classes with an invariant capsid but different tail-fiber orientations relative to the C5 axis. For 2 of the 5 classes, we obtained ∼3 Å maps of the tail region, enabling analysis of portal– capsomer interactions at the symmetry-mismatched polar vertex. This work provides the first atomic-resolution model of the complete P-SSP7 portal–tail complex. Relative to the report by Fang *et al* where tail-less T4 mutants were used, our work on intact tailed podophage shows the same portal–tail assembly occurs in multiple rotational orientations relative to an invariant capsid. We propose that P-SSP7 resolves the capsid–portal mismatch through modular attachment of the capsid shell to the portal–tail assembly in 5 rotational registers.

## 3. Results

### 3.1. Cryo-EM analysis of intact P-SSP7 resolves five distinct 3D classes

Cyanophage purification for cryo-EM remains challenging due to limited standardized, translatable protocols. We used our optimized purification workflow^23^ to generate high-quality samples for single-particle analysis. The scaled-up protocol yielded a dense suspension of P-SSP7 (>10¹² pfu/mL) that produced a near single-layer, lattice-like spread of monodisperse phages tumbling in vitreous ice (Figure S1). We collected 32,000 movies over 4 days and extracted ∼1.03 million initial particles from motion-corrected movies (Table S1). Ab-initio modeling followed by iterative 3D classification and heterogeneous refinement excluded particles lacking a tail and retained ∼660,000 tailed particles (Figure S1).

Whole-virion refinement produced a 3.02 Å map but yielded weak density for the tail region, consistent with tail heterogeneity and/or misalignment. Focused masking and 3D classification of the tail produced 5 non-redundant classes (Figure 1A, S1, and 2A) with roughly equal particle numbers (Table S2). Aligning the classes by capsid revealed 5 distinct orientations of the tail fibers, whereas aligning by fibers misaligned capsids (Figure 1A), confirming 5 unique 3D classes. At this stage, the data did not clearly indicate whether P-SSP7 had an invariant capsid with tails attached in 5 orientations or an invariant portal–tail complex with capsids attached in 5 orientations.

**Figure 1.**
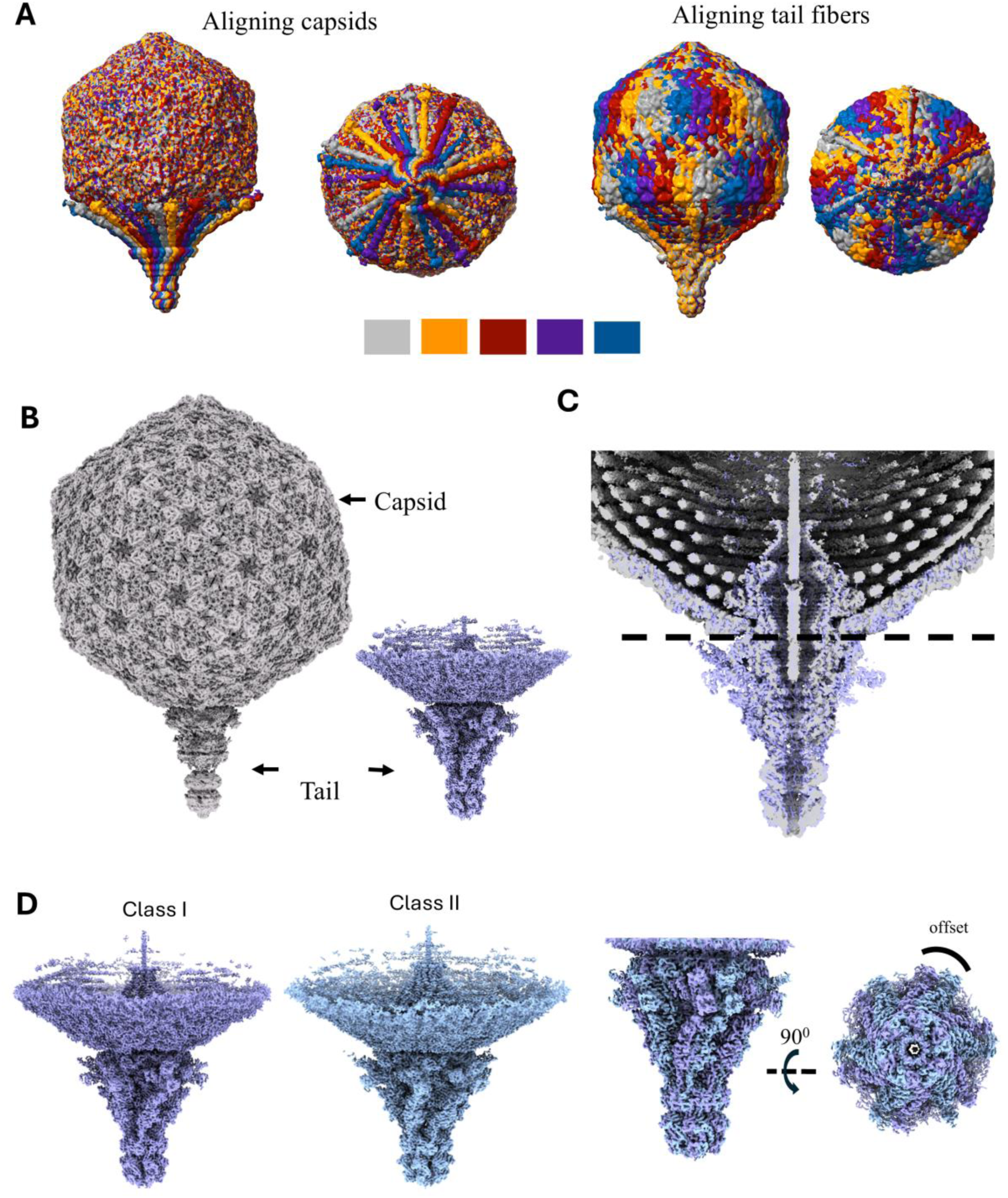
Cryo-EM map of P-SSP7 shows 5 different co-axial orientations of the tail assembly. **A)** Volumes from 3D classification, aligned relative to the capsids, show 5 different co-axial orientations of the tail fibers; volumes from the 3D classification aligned relative to the fibers show that the fibers overlap but the capsids are misaligned. B) Cryo-EM map of entire P-SSP7 (grey) phage both at 3.02 Å resolution and the map of the tail region at 3.03 Å (purple). Both maps are superimposed in (C). C) A virtual slice through the superimposed maps of the entire phage shows better resolution for the tail region in the purple map. The dotted black line indicates the region of the 12:5 symmetry mismatch. **D)** Maps of the portal-tail complex from Class I and II classes individually (purple and blue) aligned to an invariant capsid show a rotational offset relative to each other.

The shell–portal interface contains the symmetry mismatch between the unique 5-fold capsid vertex and the 12-fold adaptor/portal region (Figure 1B, dashed line). Prevailing models propose that virion assembly begins with the portal complex, proceeds through capsid assembly and genome packaging, and ends when adaptor–tail components append to the exterior of the packaged capsid. We designated particles from the 5 classes as Class I–V. Whole-virion maps for Classes I–V reached ∼3.02–3.04 Å after symmetry relaxation in I symmetry, yet each map retained low-resolution, weak nozzle density (Figure 1B,C; Figure S1). For subsequent analyses, we treated the capsid as invariant and interpreted the 5 classes as discrete tail attachment registers; under this model, the portal–tail complex rotates about the C5 axis to append to the symmetry-mismatched vertex through 5 interaction sets.

To improve tail resolution, we selected Class I and II, which showed the strongest tail density in virtual cross-sections of the full-virion maps (Figure S1, S2B). We extracted subparticles using a cylindrical mask spanning the portal complex, adaptor, fibers, and tail nozzle and including the first ring of capsomers surrounding the unique 5-fold vertex. Refinement in C1 symmetry yielded nearly identical portal–tail maps at of 3.03 Å (Class I and II) (Figure 1B–D; Figure S2), revealing high-resolution features not resolved in the full-virion reconstructions (Figure 1B,C). Refinements imposing higher symmetries (C5, C6, C10, C12) or using locally masked sub-volumes of individual components produced blurred features or lower-quality reconstructions; the whole-tail mask outperformed both strategies and supported subsequent analyses. Superposition of portal–tail maps relative to an invariant capsid revealed a rotational offset (Figure 1D), providing direct evidence for modular tail attachment at distinct rotational registers. These ∼3 Å maps represent a major improvement over previously reported nanometer-resolution maps^5^ and enable unambiguous model building.

### 3.2. Atomic modeling defines portal–tail architecture and internal interfaces

We built atomic models of portal–tail components into the Class I portal–tail map using AlphaFold predictions templated on cyanophage P-SCSP1u^20^ (EMDB-35174; PDB 8I4M, 8I4L). We iteratively refined the portal, adaptor, fiber, and nozzle models against cryo-EM density to generate a complete P-SSP7 portal–tail model with overall architecture analogous to P-SCSP1u (Figure 2A). We also fitted portal, adaptor, and nozzle models into the Class II map; unless noted, we describe Class I.

**Figure 2.**
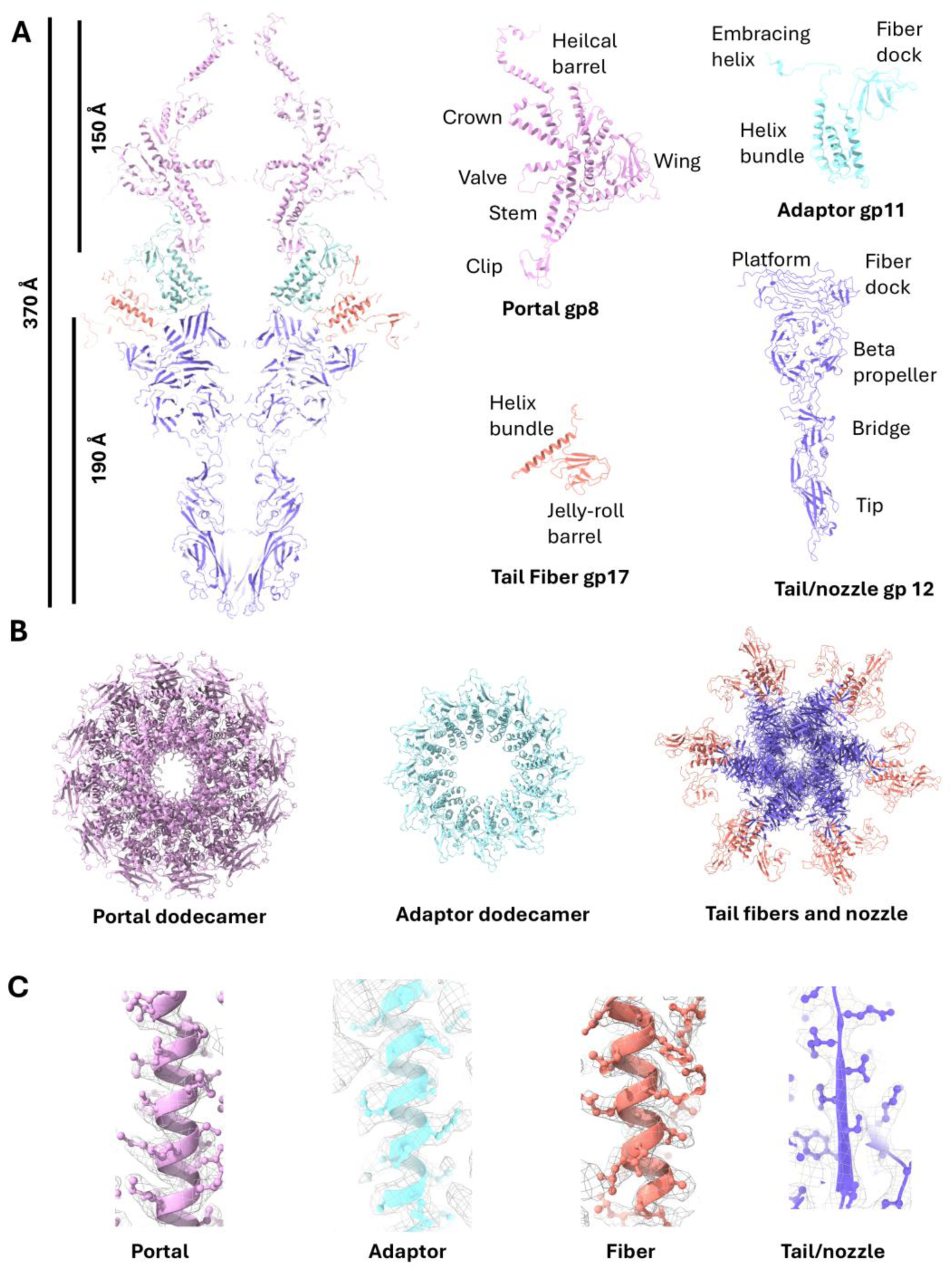
Structure of the portal and tail complex of P-SSP7. **A)** Virtual slice through the model of the entire portal and tail complex with lengths of the entire complex and of just the portal and nozzle assemblies. The panel also shows domain-level organization of the monomers of the portal (pink), adaptor (cyan), fibers (brown), and tail/nozzle (purple). **B)** Individual models of the portal, adaptor, tail fiber, and nozzle with domains annotated following the convention for other cyanophages with similar architecture^20,37,53^. **C)** Representative regions of the model–density fit for each subunit, shown as atomic models (colored ribbons with side chains) within the experimental cryo-EM density (grey mesh): portal (pink), adaptor (cyan), fiber (brown), and tail/nozzle (purple).

#### Portal

The portal complex comprises a gp8 dodecamer (522 aa). We modeled residues 1-502; density did not resolve the C-terminal loop of 20 residues. The portal contains crown, stem, and clip regions oriented toward the central pore that houses and interacts with the genome. The wing domain faces outward and adopts an α/β fold with a C-terminal helical barrel. The portal shows C12 symmetry approximately around the C5 axis; the helical barrel and the valve region form 2 constrictions that interact with the genome (Figure 2A,B, pink). Relative to P-SCSP1u, gp8 shows three differences: the N-terminal region lacks a disordered loop present in P-SCSP1u; the wing contains a disordered loop (aa 177–197) lacking density; and the C-terminal helical barrel is more steeply angled and shorter. The region spanning aa 345–367 which putatively forms a valve within the genome-ejection pore, also lacked corresponding density. The portal measures 150 Å in length and 170 Å in diameter.

#### Adaptor

Beneath the portal, a cylindrical gp11 dodecamer (204 aa) forms the adaptor (Figure 2A,B, cyan). Gp11 seals the capsid to the tail nozzle and provides attachment points for tail fibers. Each gp11 contains an embracing helix that contacts gp8, a 3-helix bundle with a hydrophobic core that contacts the tail, and a β-sheet roll that provides a surface for fiber docking. The adaptor fold matches adaptor proteins from other tailed bacteriophages, including P-SCSP1u^20^.

#### Tail/nozzle

The nozzle comprises a gp12 hexamer with overall organization similar to P-SCSP1u. Gp12 includes a platform domain attached to the adaptor, a fiber dock, a β-propeller region, and a tip housing receptor-binding residues (Figure 2A,B, purple). β-sheets dominate the structure, with 6 gp12 monomers forming an overall funnel-shaped organization. Gp12 is 175 aa longer and has a more elongated domain organization than the P-SCSP1u nozzle monomer (Figure S3A). Multiple sequence alignments of nozzle proteins from related cyanophages identified an additional domain (aa 176–340) of ∼170 amino acids absent in P-SCSP1u (Figure S3B). This additional domain interacts with host receptors and makes the P-SSP7 nozzle ∼50 Å longer than that of P-SCSP1u. Although AlphaFold predicted the full nozzle, the predicted architecture deviated from cryo-EM density (Figure S3C), requiring map-guided fitting and refinement. The portal–tail assembly measures 370 Å, ∼40 Å longer than the corresponding assembly in P-SCSP1u, primarily due to the extra gp12 domain.

#### Fibers

We fitted the N-terminal 117 aa of tail fibers into density contiguous with the adaptor and nozzle. Three gp17 monomers assemble into a single trimeric fiber in a concentric propeller configuration. Distal domains remained visible only as weak density and could not be resolved. Each monomer contains a helix bundle that attaches to the nozzle and a jelly-roll domain that attaches to the adaptor (Figure 2A,B, brown).

#### Unique interactions within the portal–tail complex

From portal to nozzle, a minimal set of 2 gp8 chains (P1 and P2), 2 gp11 chains (A1 and A2), 1 gp17 trimer (F1, F2, and F3), and 1 gp12 monomer (T1) captured all unique interactions within the portal–tail complex (Figure 3). We used PISA^24^ to quantify interfaces and map all the unique inter- and intra-component interactions within the portal-tail complex. The two gp8 chains formed a strong interface with 48 H-bonds, 5 salt bridges, and a shared surface area of ∼4808 Å². The gp8 stem interacted with C-terminal residues of gp11 at the embracing helix (P1:P49-A1:R204) via extensive hydrogen bonding, and the gp8 clip formed a salt bridge with flexible gp11 loops (P2:K295-A2:E54) (Figure 3A,B). The terminal Arg of gp11 contacted residues from 2 portal chains (Figure 3B). Gp11 monomers formed a stable dodecamer via extensive hydrogen bonds, salt bridges, and a buried surface area of ∼2140 Å2 between two monomers. The adaptor ring sat at the interface of the capsid and solvent-exposed exterior and contacted the nozzle platform through loops in the triple-helix bundle (Figure 3C). Every alternate gp11 chain contacted 1 gp12 chain (Figure 3A,C), securing portal, adaptor, and nozzle.

**Figure 3.**
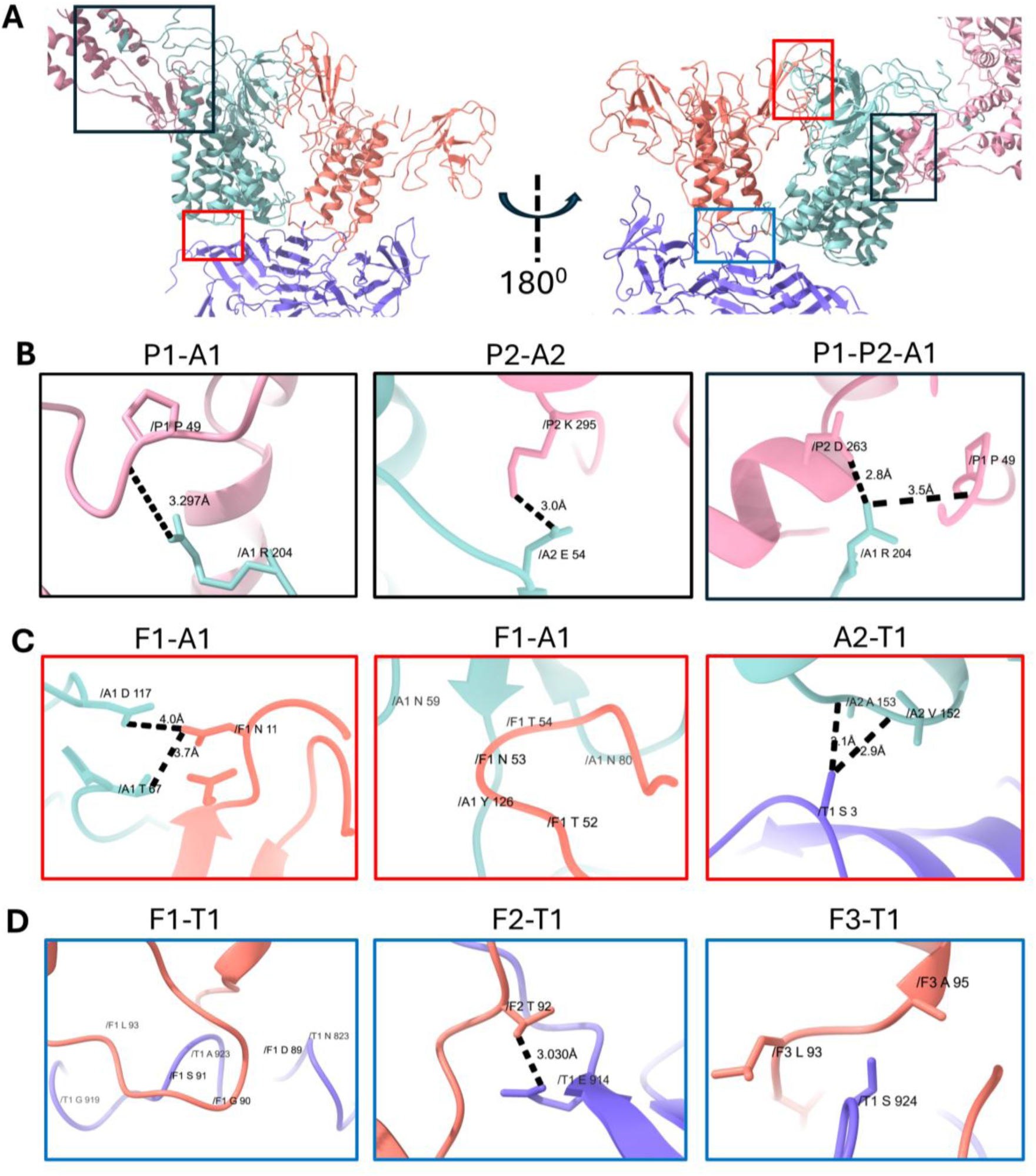
Interactions in the portal-tail complex. **A)** Zoomed-out views of 2 chains of the portal complex (P1, P2), 2 chains of the adaptor (A1, A2), a fiber trimer (F1, F2, F3), and 1 chain of the nozzle/tail (T1) from two angles (180° rotation). Regions bounded by black, red, and blue boundaries are used to zoom in to show representative interactions between side chains of the different chains in the portal-tail complex. **B)** R204 from A1 interacts with P49 from P1 and D263 from P2. E54 from A2 also forms a salt bridge with K295 from P2. **C)** H-bonds formed by A1 with F1 and a longer loop interacting with A1. A2 forms H-bonds with T1. **D)** H-bonds between fiber monomers and the fiber-docking domain of T1, shown for each F–T interface. **F1–T1**: loop residues F1 D89, F1 G90, F1 S91, and F1 L93 contact T1 N823, T1 G919, and T1 A923. **F2–T1**: F2 T92 T1 E914. **F3–T1**: F3 L93 and F3 A95 are positioned near T1 S924.

Each fiber formed a gp17 trimer via N-terminal residues. Two gp17 monomers (F1 and F2) contacted two adjacent gp11 monomers (A1 and A2) (Figure 3A,C). The F1–A1 interface was especially extensive, with a 909 Å^2^ buried surface area at the jelly-roll domain (Figure 3C). The F2–A2 interface remained localized to A2:Thr28 and the loop connecting the jelly-roll domain to the α-helix on F2. In addition, all 3 loops (aa 90-95) of a gp17 trimer near the helix bundle attached to the gp12 anchor domain (aa 823 and 919-925) (Figure 3D). The jelly-roll domain of the third gp17 monomer remained solvent exposed. The N-terminal ∼100 residues of the fiber trimer therefore function as a stabilizing plug by linking 2 gp11 monomers and 1 gp12 monomer. Gp17 did not contact gp8.

### 3.3. Modular attachment resolves the 12:5 mismatch at the unique vertex

The portal–tail complex is surrounded by a ring of 5 capsid hexons of the shell protein gp10 (chains a-f) at the unique 5-fold vertex (Figure S4A,B). We fitted gp10 models for hexon 1 (chains a1-f1) into the map (Figure S4D). Gp10 adopts an HK97-like fold with the characteristic N-arm, A domain, E loop, F loop, and P domain^25^. Chains a1 and c1-f1 fit density well, but density did not resolve the N-arm of chain b1 (aa 1-30), indicating complete disorder despite juxtaposition to the gp8 dodecamer (Figure S4C-E). Only chains a and b from each hexon contacted gp8.

To analyze portal–capsomer interactions, we built models of gp10 chains a and b from all 5 hexons in Class I and II maps (a1b1 to a5b5) (Table S3-S5). We removed segments lacking density: the disordered gp8 loop aa 177-197 and the gp10 chain-b N-terminal 30 residues in all 5 hexons in both classes. Aligning and superimposing gp8(P1-P12):gp10(a1-a5) between classes showed perfect alignment of gp8 but a ∼12° rotational offset of gp10 (Figure 4A,B; Figure S5A,B). Gp10 a/b chains interacted primarily with the N-terminal 45 residues of gp8, which form a helix and adjacent loop in the wing domain (Figure 4B–D, green).

**Figure 4.**
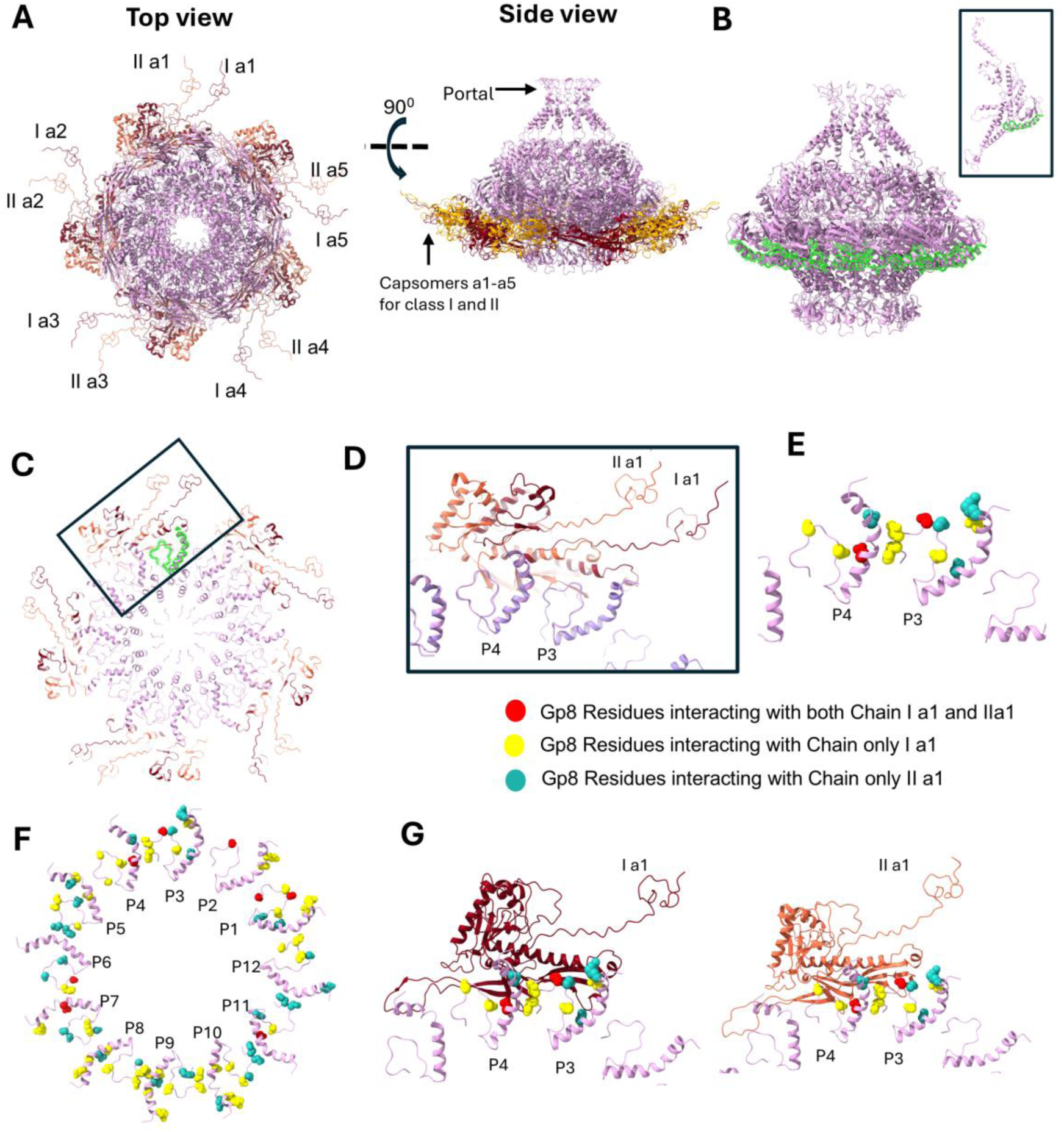
Interactions of the portal and capsomers from 2 classes of tails. **A)** Ensemble of the gp8 and gp10 chain a from each of the 5 hexamers at the unique 5-fold vertex from Class I and II of tail assemblies. Top view is along the C5 axis and side view orthogonal to the C5 axis. Chains a1-a5 for Class I (dark brown) and Class II (orange) are slightly offset from each other (hence forth called I a1-a5 and II a1-a5). **B)** The capsomers interact with the N-term residues 1-45 of the portal protein gp8 (highlighted in green). The inset shows the location of residues 1-45 on a single gp8 chain. **C)** Virtual slice along the C5 axis of the ensemble showing gp8 1-45 (highlighted) and interacting with chain a1 from Class I and II in the black boundary. **D)** A zoomed in view of the region in black boundary in (C) shows while the Chain a1 form equivalent capsomers slightly off set from each other and interact with chains P3 and P4 of gp8. Pink and purple colors for P3 and P4 are from the gp8 chains from 2 classes of tail assemblies perfectly aligning to each other. **E)** Residues interacting with I a1 and II a1 on gp8 chains P3 and P4 shown with legend. **F)** Residues interaction with Chains I a1-a5 and II a1-a5 mapped on to the N term region of gp8 dodecamer **G)** Comparative mapping of interactions of Ia1 and II a1 on gp8 P3 and P4 showing redundant and unique interactions.

We quantified interactions between gp8(P1-P12) and gp10 a/b chains across all 5 hexons in both maps using PISA (Figure 4; Figure S5). The analysis identified three interaction classes: interactions unique to Class I or Class II, interactions shared across both classes, and interactions in which conserved gp8 residues contacted gp10 in both classes but paired with different gp10 residues (Figure 4E–G; Figure S5; Table S5). The rotational offset observed in maps (Figure 1D; Figure 4D–F) also appeared in the gp8:gp10-chain-a1 interfaces (Table S6). For example, gp8-P3 interacted only with gp10-a1 in Class I but interacted with gp10-b5 and gp10-a1 in Class II (gp10-b5 originates from an adjacent hexon; not shown). Projection onto the gp8 N-terminal loop showed that residues interacting with gp10 chain a in Class II (yellow) shifted relative to those interacting with gp10 chain a in Class I (light sea green) (Figure 4E,G; Figure S5C-E).

Ser and Asp residues on gp8 mediated ∼50% of gp8–gp10 interactions; 9/12 gp8 chains interacted with gp10 through a Serine. Conversely, gp10 residues between aa 325-335 contributed ∼50% of gp8 interactions, though not necessarily via Ser residues (Table S5). Several gp8 residues participated in interactions in both classes but formed bonds with different gp10 residues (Table S5 highlighted). For example, P1:Ser39 interacted with a5:Arg48 and a5:Asp50 in Class I but interacted with a5:Val330 in Class II. We also identified gp8 residues that interacted uniquely with gp10 residues either only in Class I or only in Class II (Figure 4E,F; Figure S5C-E; Table S5).

## Discussion

The major finding of this study is the 5 distinct 3D classes in which the same portal–tail assembly attaches to an invariant capsid in different rotational orientations. We report atomic-resolution maps of 2 of the 5 classes of the P-SSP7 portal–tail complex at 3.03 Å resolution each. Previous work reported only domain-level organization using segmented tail components^5^. In addition to improvements in cryo-EM instrumentation and software since 2010, two factors enabled the structural detail reported here: high-quality sample preparation^23^ and a dataset ∼10× larger than those used for cryo-EM studies of other podoviruses^5,18–20^ (Table S1). This dataset enabled refinement using ∼100,000 boxed-out subparticles per class for capsids and portal–tail assemblies. Asymmetric C1 reconstructions preserved class-specific features that would be washed out by imposing higher symmetry. Consistent with this, Liu *et al.* used 36,000 particles in the final reconstruction, compared with ∼100,000 particles per class here, which likely contributed to earlier domain-level tail resolution.

The gp10 model built into our higher-resolution map showed no major differences relative to published capsid structures. Although we initially interpreted the classes as 5 portal–tail orientations attached to an invariant capsid, our gp8:gp10 analysis aligned gp8 between Class I and II, consistent with an invariant portal–tail assembly and distinct capsid rotational registers. Fang *et al*.^19^ reported an analogous phenomenon using a tail-less T4 mutant, identifying 5 capsid classes slightly rotated about the C5 axis and proposing that portal conformational changes compensate for the symmetry mismatch at the unique 5-fold vertex. In P-SCSP1u, reports suggest that the portal at the 12:5 interface loses strict 12-fold symmetry while surrounding capsomers remain stable.

Several distinctions separate our study from reports by Fang *et al* and Cai *et al*. The T4 analysis used a tail-less mutant; T4 remains challenging for cryo-EM, and our current understanding relies on structural studies employing components of T4 expressed *in vitro* or ingenious use of structural mutants^26–29^. In contrast, we analyzed intact wild-type P-SSP7 virions. In addition, maps publicly available from Fang *et al*. (EMD-20960 and 20961) applied C12 or C5 symmetry, whereas we reconstructed the P-SSP7 portal–tail assembly in C1 symmetry, consistent with studies in which C1 reconstructions reveal features otherwise lost after applying higher-order symmetry^12,16,30,31^. Notably, P-SCSP1u was not reported to show modular tail arrangements despite genetic and structural similarity to P-SSP7, and Cai *et al*. provided only domain-level information on portal–capsomer interactions.

Our understanding of podovirus assembly derives largely from extensive studies of P22 and T7 podophages^17,32–34^. In these models, the portal complex assembles first, capsomers then assemble the capsid (sometimes aided by cement proteins), headful genome packaging completes the head, adaptor proteins seal the capsid, and tail components assemble externally; tail structures therefore grow outward from the portal ring during assembly^35^. Genome structure analyses suggest that P-SSP7 follows a T7-like assembly model^11^. Podovirus tails show extreme diversity^13,36,37^ and Syn5, P-SSP7, and P-SCSP1u differ in tail size. P-SSP7 has a longer tail with an extra bridging domain, and P-SCSP1u may represent a derived state in which a fragment of the tail protein was deleted. Tail-length variation even among related cyanopodoviruses suggests that these viruses evolved diverse tail machineries to attach, infect, and inject genomes into the same host.

We propose that the portal dodecamer and capsid shell adopt 5 distinct rotational registers at the unique 5-fold vertex, yielding (1) redundant and class-specific portal–capsomer interactions and (2) adaptor/nozzle assembly that follows portal orientation rather than shell orientation. We found all 5 orientations at roughly equal abundance (∼20% each). Attempts to classify into >5 classes led to convergence into 5 unique classes, indicating uniform distribution and no dominant register. We estimate that each class differs by ∼12° based on inter-class fiber offsets and that a hypothetical Class VI would converge with Class I. We propose that modular attachment with 5 unique registers resolves the 12:5 symmetry mismatch in P-SSP7 and represents the first reported instance of such a mechanism. The 2 tail apparatus conformations in P22 and SF6 are rotationally offset by 6^0^ relative to the 12° offset for P-SSP7^21,22^.

Placing our findings in the context of prior work reveals that dsDNA phages employ a spectrum of mechanisms to address the portal-capsid symmetry mismatch. In bacteriophage T7^18^, the internal packaging apparatus adopts uncorrelated, tandem symmetry mismatches where components do not need to maintain fixed rotational correlation relative to the capsid. In HK97, scaffold flexibility can accommodate a 12-fold pore at the 5-fold vertex^16^, and in phi29 direct most recent Cryo-EM reports support local oscillation or rotation rather than large scale connector rotation during packaging, partially supporting rotary mismatch-resolution models^17,38^. Continuous conformational morphing at the mismatched vertex, as reported for a tail-less T4 mutant^19^, represents yet another solution. Alternatively, recent Cryo-EM studies show that the tail apparatus in the model podophage P22 and a related coliphage SF6 exists in two equivalent conformations ^21,22^, or energetic minima, which accommodates the symmetry mismatch. Against this backdrop, P-SSP7 is distinctive: rather than a single fixed geometry or continuous morphing, the intact wild-type virion populates five discrete, equally abundant (∼20%) rotational registers separated by ∼12°, each defined by redundant and class-specific gp8–gp10 interfaces. This quantized modularity, resolved here in intact particles by C1 reconstruction, suggests that P-SSP7 accommodates the 12:5 mismatch through a defined set of interchangeable interaction sets rather than through symmetry-averaged plasticity, extending the conceptual range of mismatch-resolution strategies in tailed phages.

Future studies could also investigate if tail apparatus from other known or novel phages has multiple tail conformations and if there are phages that display 3 or 4 different conformations in addition to the currently known examples. Alternatively, mutants that enrich a single register could be generated to test whether register identity alters infection dynamics. Cell-free protein expression can produce fully assembled, infectious phages^39^ and could probe early P-SSP7 assembly by expressing gp8 and gp10 separately, together, and sequentially. If portal assembly nucleates capsid formation, such approaches could yield capsids assembled around an invariant portal in 5 orientations, enabling cryo-EM analysis of assembly intermediates.

Based on our current knowledge no other podophage shows this modular arrangement with 5 registers. However, multiple tail conformations with rotational offsets in some multiple of 6^0^, observed across viruses from different families (P-SSP7, T4, SF6 and P22), could be a mechanism of resolving the 12:5 symmetry mismatch at the unique polar vertex more widely adopted by phages. Intact T4 and other tailed phages may also adopt modular portal–tail orientations relative to an invariant capsid as a general mechanism to resolve the 12:5 mismatch. Although direct biochemical evidence for P-SSP7 assembly intermediates remains limited, transcriptomic studies—an initial microarray study^8^ and our RNA-seq work^10^—show that portal (gp8) and scaffolding (gp9) genes express earlier than the major capsid protein gene (gp10). This temporal separation supports portal-first nucleation models established for P22^40,41^ and conserved across tailed dsDNA phages and herpesviruses^42^. Nevertheless, cryo-EM alone cannot determine whether an invariant capsid accommodates 5 portal–tail orientations or whether 5 capsid registers form around an invariant portal–tail complex. We therefore define the 5 classes as distinct modular arrangements of the shell and portal–tail complex.

## Online Methods

### Cyanobacteria Host Culture and Cyanophage

*Prochlorococcus marinus* strain MED4 (CCMP1986) was obtained from the Provasoli-Guillard National Center for Marine Algae and Microbiota (NCMA, East Boothbay, ME), and phage P-SSP7 was provided by the Chisholm Laboratory (Massachusetts Institute of Technology). MED4 was cultured under a 12-h light:12-h dark cycle (25 μmol quanta m⁻²⋅s⁻¹) at 21 °C in natural seawater-based Pro99 medium. Cultures were maintained in volumes ranging from 5 to 100 mL in sterile test tubes and cell culture flasks. For routine P-SSP7 propagation, exponentially growing MED4 cultures were infected and harvested after complete lysis (6–10 days post-infection).

### Large Scale MED4 Host Cultures and Phage Production (up to 40 L)

#### Scale-up of MED4 Cultivation and Infection

Detailed protocols for scaling MED4 culture, infection, and P-SSP7 purification are provided in our previous work^23^. Briefly, *Prochlorococcus marinus* strain MED4 was adapted to local Salish Sea seawater over a two-month period using 20% incremental increases. Growth medium was supplemented with 2× Pro99 N and P concentrations and 1× trace metals to achieve stable growth patterns and higher biomass density. Cultures were maintained under a 12-h light:12-h dark cycle (35 μmol quanta m⁻²⋅s⁻¹) at 21 °C with gentle aeration for CO₂ supply and mixing. For final scale-up, MED4 was cultivated in 12 L food-grade PET carboys, with aeration provided through foam stoppers with 0.2 µm filters for sterile gas exchange. Two to four carboys (10 L each) of mid-exponential phase culture (0.1–1 × 10⁸ cells/mL) were infected with freshly prepared P-SSP7 lysate at an MOI of 0.5–1 × 10⁻². Following a 60-minute adsorption period, cultures were incubated under 24-h continuous light until complete lysis (6–10 days), with supplemental nutrients added every 3–4 days to support host metabolism during infection.

#### Phage Harvesting and Purification

Following complete lysis, lysates were pre-treated with 0.2 U/mL DNase I for 30 min at room temperature to reduce viscosity, followed by NaCl addition (116 g/L; 2 M final concentration) with gentle stirring and 60-min incubation at 4 °C to facilitate release of phage particles from membrane vesicles and cellular debris. Cellular debris was then removed by centrifugation (26,000 × g, 40 min, 4 °C). The optimized purification protocol included sequentially: (i) PEG 8,000 precipitation (10% w/v with 2 M NaCl), (ii) ultracentrifugation through a 38% sucrose cushion (113,000 × g, 3.5 h, 4 °C), and (iii) two-step CsCl density gradient separation: a first gradient at 1.2–1.65 g/mL (154,000 × g, 3.5 h, 4 °C) followed by a second step (98,000 × g, 12–16 h, 4 °C) for enhanced phage purity. The visible phage band was collected and dialyzed through a stepwise procedure to gradually remove CsCl. Purified phages were stored at 4 °C for subsequent applications. Detailed protocols for phage production, harvesting, and purification are provided in^23^.

### Quantification of cyanobacteria and phage

#### Quantification of Host Cells and Phage Particles

Host cell density and total phage particle concentrations were quantified using routine methods established in our previous work (Bohutskyi et al. 2026). MED4 cell density was monitored by optical density at 750 nm using a Genesys 20 Spectrophotometer (Thermo Fisher Scientific); OD₇₅₀ values were converted to absolute cell counts using a previously established flow cytometry-based calibration curve (R² = 0.935, p < 0.001). Total phage particle concentrations were determined by epifluorescence microscopy following DNase I treatment, SYBR Gold staining, and filtration onto 0.02 μm pore size Anodisc filters (Cytiva Whatman, 09-926-34). Samples were imaged using a Leica epifluorescence microscope (100× oil objective, 488 nm excitation, 525 nm emission) and particles counted by automated thresholding from ∼20 fields per sample in Fiji (ImageJ) ^43^.

#### Infectious Phage Titer by MPN Assay

To determine the concentration of infective phage particles, a Most Probable Number (MPN) assay was performed in 96-well microtiter plates as described in our previous work^23^. Serial 10-fold dilutions of phage samples (10⁻³ to 10⁻¹⁷) were prepared in Pro99 medium (2× N and P). For each dilution, 20 μL was added to wells containing 30 μL of exponentially growing MED4 cells (0.1– 1 × 10⁸ cells/mL). After a 1-hour adsorption period under standard growth conditions (35 μmol quanta m⁻²⋅s⁻¹, 21 °C), 200 μL of Pro99 medium was added to each well to bring the total volume to 250 μL. Plates were sealed and incubated for 1–2 weeks, with chlorophyll fluorescence monitored every 2–3 days using a Biotek Cytation C10 plate reader (485 nm excitation, 675 nm emission); wells showing significant reduction in fluorescence relative to controls were scored as positive for host lysis. Six replicates were performed per dilution, and MPN estimates with confidence intervals were calculated using the MPN R package. ^44,45^

### Cryo-EM grid preparation and data collection

3 µL of PSSP-7 suspension was loaded on to glow discharged Quantifoil grids (R 1.2/1.3 or R 2/2, 300 mesh). Grids were blotted for 2.5–3.5 s and plunge-frozen in liquid ethane using a Leica EM GP2, then stored in liquid nitrogen until use. For screening and data collection, grids were loaded onto a 300 keV Titan Krios G3i (Thermo Fisher) equipped with a K3 direct electron detector and a BioContinuum energy filter (Gatan Inc.) with a 20 eV slit. Datasets were collected using SerialEM at 81,000× nominal magnification in counted mode (calibrated pixel size 1.1 Å, F-cropped to 1.501 Å), with a total dose of ∼45 e⁻/Å² across 46 subframes and a defocus range of−0.75 to −1.5 µm.

### Image Processing

All movies were processed using cryoSPARC Live and cryoSPARC ^46^. Motion correction and CTF estimation were performed using default parameters, and initial particle extraction used the built-in *blob picker* with a box size of 1,200×1,200 pixels ^47^. Details about particle numbers at each step are listed in **Supplementary Table 1, 2** and **Figure S1**. Initial subsets of particles were subjected to reference-free 2D classification before discrete and diverse classes were chosen to re-extract particles using template picking. Multiple rounds of classification were performed to exclude junk and non-homogenous classes. Ab-initio models were generated using a subset of these particles and C1 symmetry. Heterogeneous refinement was used to exclude 40% of the particles without the tail assembly. The remaining 60 % intact particles were further refined in 3D against ab-initio models using icosahedral symmetry with symmetry relaxation. A total of 669,000 particles were used to obtain a cryo-EM map of the virus at a resolution of 3.02 Å. Per-particle local CTF refinement was performed before the final round of homogeneous refinement^48^.

3D classification resulted in 5 distinct classes in which the capsids aligned perfectly; however, the tail fibers had 5 different coaxial orientations. Each of the 5 classes was then processed further individually with symmetry relaxation and a 3.02 Å map was obtained for the entire phage. Custom cylindrical masks of 500×300 pixels were designed and centered on a region of the virus containing the entire tail, portal complex, fibers, and the capsomers interacting with the tail around the unique 5-fold vertex (referred to here as the *tail region*). The masks were used to extract sub-particles of 500×500 pixels from the motion-corrected micrographs. These subparticles were used to reconstruct the tail region from 2/5 classes in C1 symmetry. Cryo-EM maps of the tail regions from 2/5 classes were obtained at 3.03 Å resolution each. All resolutions were estimated using the gold-standard 0.143 FSC criterion, and maps were visualized using UCSF ChimeraX^49^.

### AlphaFold, Modelling and refinement

Cryo-EM maps of the capsid and the tail region were z-flipped before model fitting. Initial models for the portal protein 12mer (gp8), adaptor 12mer (gp11), fibers (gp17) and nozzle/tail 6mer (gp12) were obtained using AlphaFold with PDB:8I4M as a reference. PDB:2X8D was used as the reference template for the capsid protein (gp10). AlphaFold was also used to obtain the model for the entire assembly of the tail complex prior to fitting. Models of all individual components were docked into the map of the tail region, and smaller sub-volumes were extracted using the MapBox feature in Phenix. Model fitting was iteratively improved using Phenix and Coot^50–52^.

## Author contributions

A.D.P., P.B., M.S.C., J.E.E. designed the research.

A.D.P., P.B., N.S performed wet lab experiments

A.D.P. and J.E.E performed data processing and analysis

A.D.P and P.B wrote the first draft

M.S.C., J.E.E. provided supervision, mentorship, funding and resources.

All authors contributed to editing and writing the paper.

## Acknowledgements

The research (NW-BRaVE) described in this paper is supported by the BRaVE project funded by the U. S. Department of Energy (DOE), Office of Science, Office of Biological and Environmental Research, under FWP 81832. A portion of this research was performed on a project award (61054) from the Environmental Molecular Sciences Laboratory, a DOE Office of Science User Facility sponsored by the Biological and Environmental Research program under Contract No. DE-AC05-76RL01830. Pacific Northwest National Laboratory is a multi-program national laboratory operated by Battelle for the DOE under Contract DE-AC05-76RL01830. A portion of this research was supported by NIH grant R24GM154185 and performed at the Pacific Northwest Center for Cryo-EM.

## Data availability

The maps for the entire virion (EMD-78075); Class I portal complex and its model (EMD-78014) and models for the tail complex (37AN) and shell protein (37AT); and Class I portal complex and its model (EMD-78027) and models for the tail complex (37CZ) and shell protein (37AB) have been deposited to EMDB and will be released upon publication.

