## Supplementary_Information for "Five Rotational Registers Define Modular Portal-Capsid Assembly in Cyanophage P-SSP7"

<u>Content</u>

Figures S1-S5

Tables S1-S6

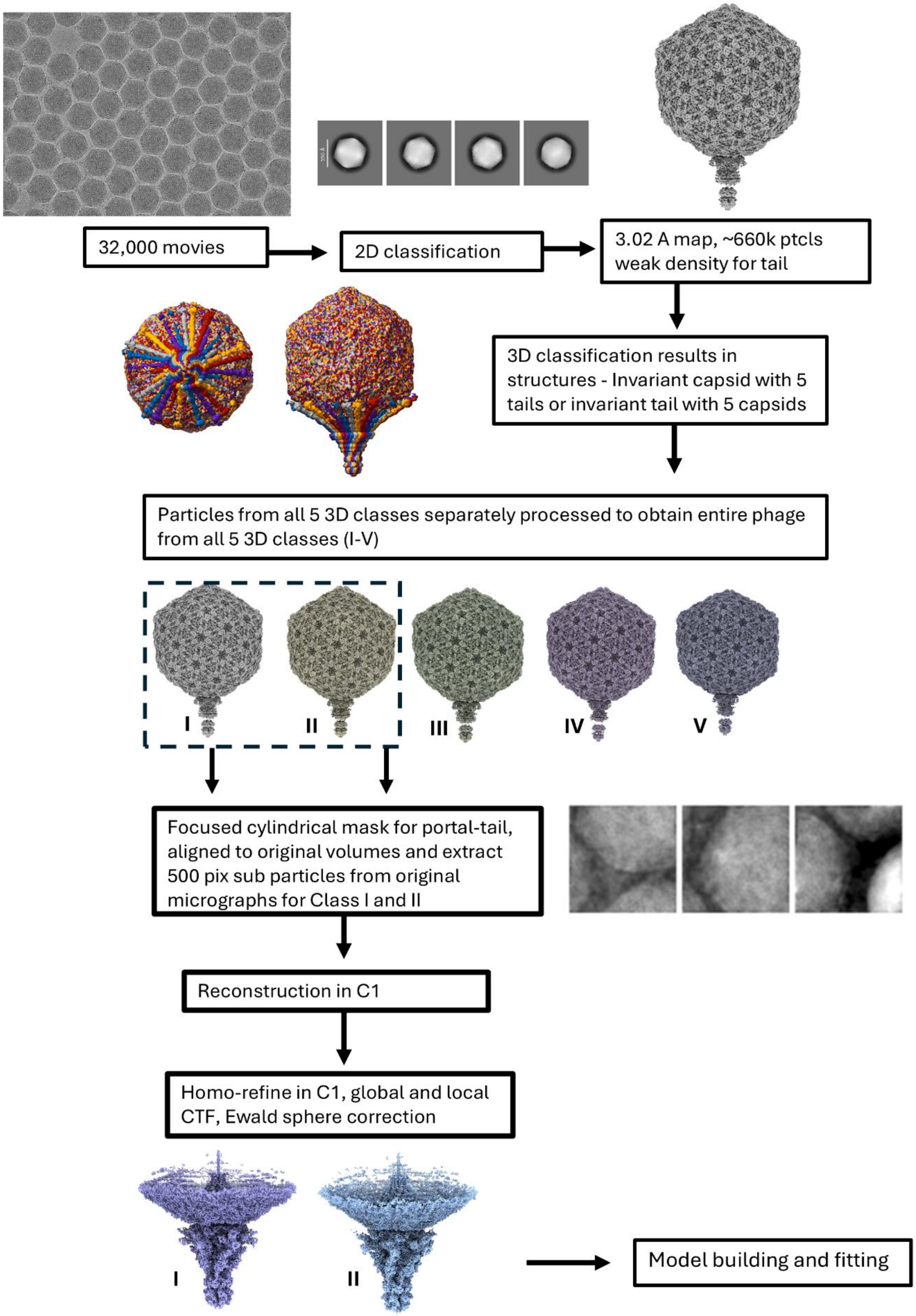

**Figure S1.** Image processing workflow for obtaining cryo-EM for the entire virions from the 5 different 3-D classes (Classes I-V) and portal-tail complex from Class I and II assemblies.

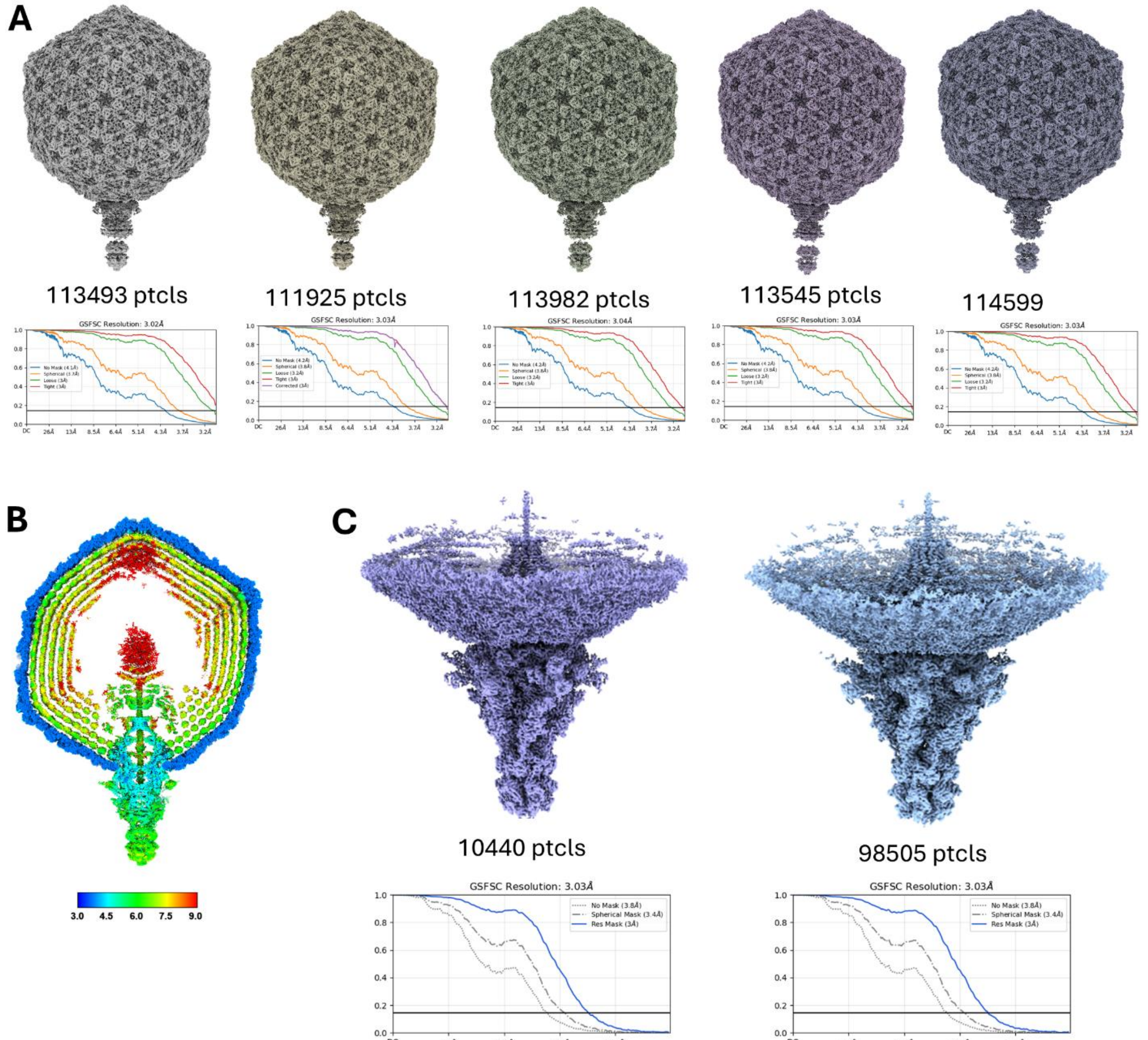

**Figure S2 Cryo-EM maps obtained from analysis of the 5 3D classes of P-SSP7**. A) Cryo-EM maps of the entire P-PSSP7 virion from Classes I-V at 3.02-3.04 Å resolution based on 0.143 FSC gold standard. B) A virtual slice through the resolution heat map of the from Class 1 showing the capsid shell resolved at ~3 Å resolution and regions of the portal tail complex resolved at 4.5-7.5 Å resolution. C) Cryo-EM maps of the portal-tail region from Class I and II with resolution at 0.143 FSC.

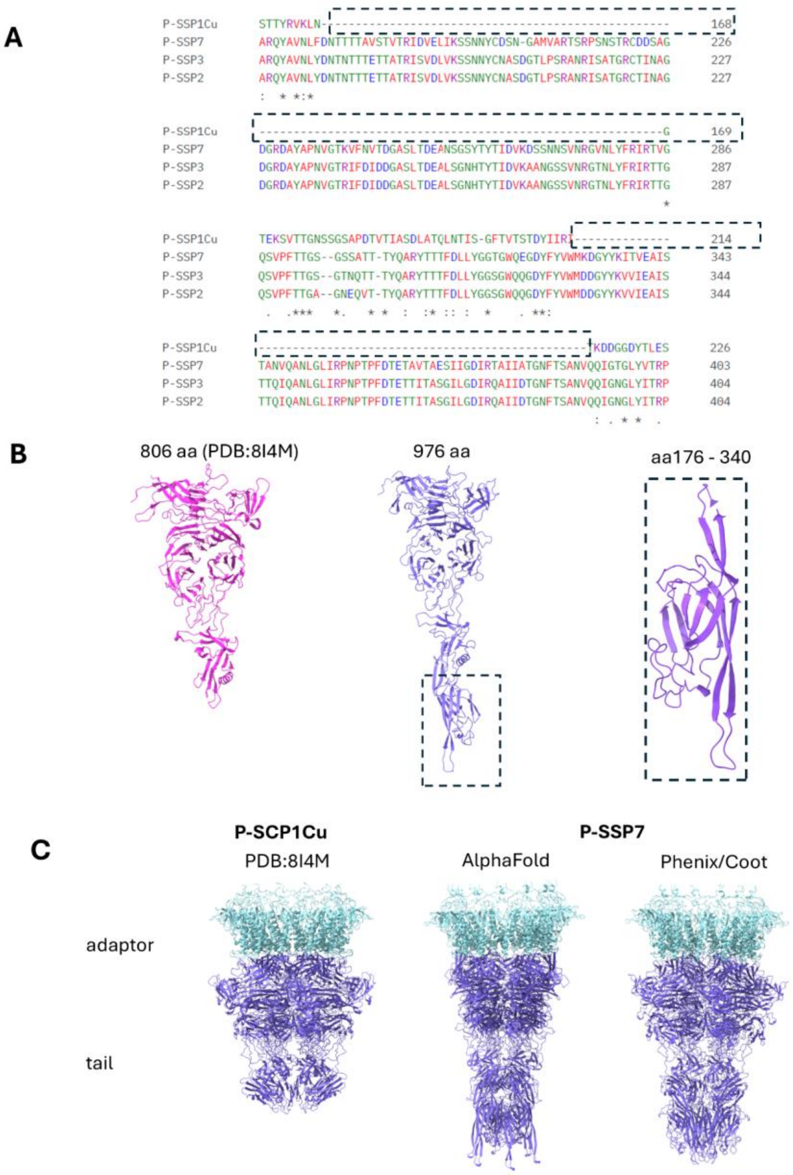

**Figure S3 –Comparing structure of the tail/nozzle between P-SSPP7 and related podophage PSCP1-Cu.** A) Multiple Sequence alignment for the nozzle protein between P_SCP1Cu and P-SSP2, 3 and 7 shows 2 additions in the primary sequence (dotted boundary) that are not present in the more basal P-SSP1Cu. B) Comparing the monomer of the nozzle protein between P-SCP1Cu (pink) and P-SSP7 (purple). The monomer has an extended structure with an additional domain between aa 176-340 which now contains the membrane binding residues. The homolog of the "tip" domain in this structure becomes a bridging domain which connects the tail to the B-propeller domain. C) Part of the tail complex containing the adaptor (cyan) and nozzle (purple) is compared between the published model for P-SCP1Cu and P-SSP7 AlphaFold and Phenix and Coot fitted and refined final model.

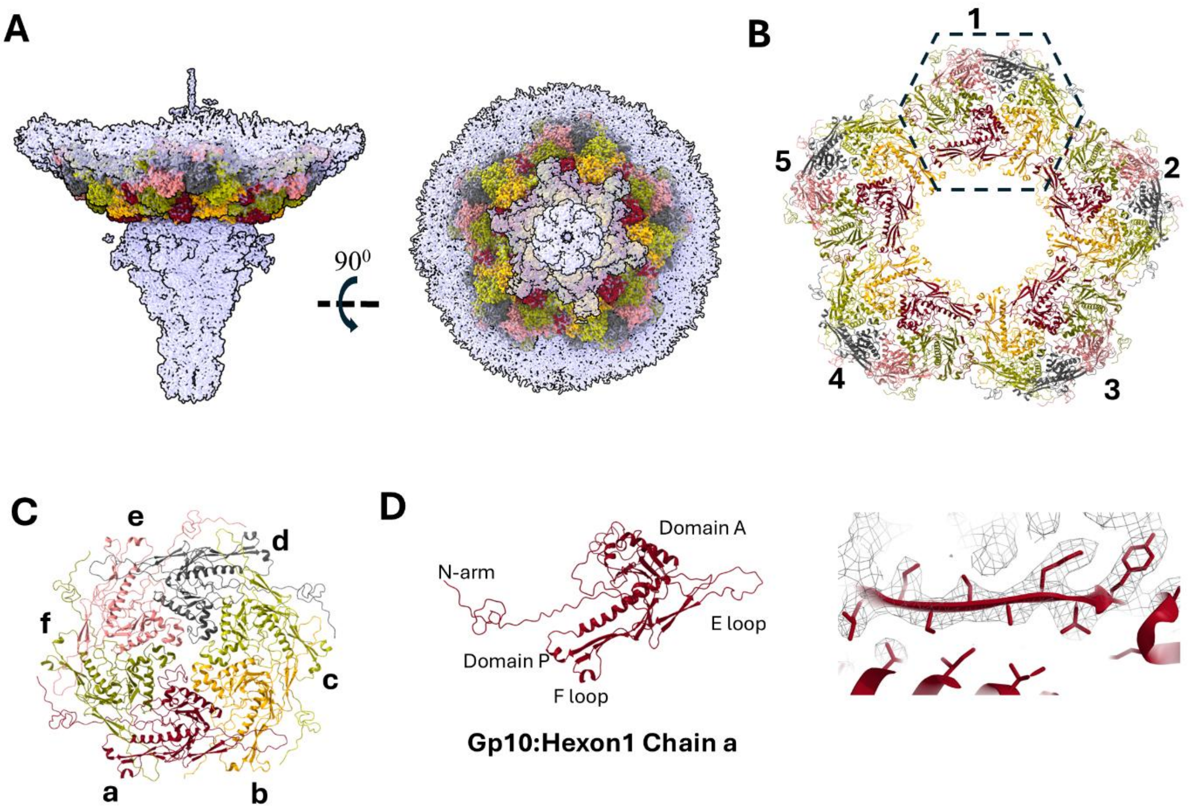

**Figure S4 Structure of the shell protein gp10 at the unique 5-fold vertex of the hexon of P-SSP7.** A) Models of models of gp10 hexons at the unique 5-fold vertex docked into the Class I portal-tail region map. B) 5 hexons at the unique vertex with Hexon 1 refined. C) Zoomed in view of Hexon 1 indicated in the dashed boundary showing 6 chains of gp10 (a-f). D) Model of Chain a from Hexon 1 showing the HK97 fold and a zoomed in view showing fitting of the model in the EM density.

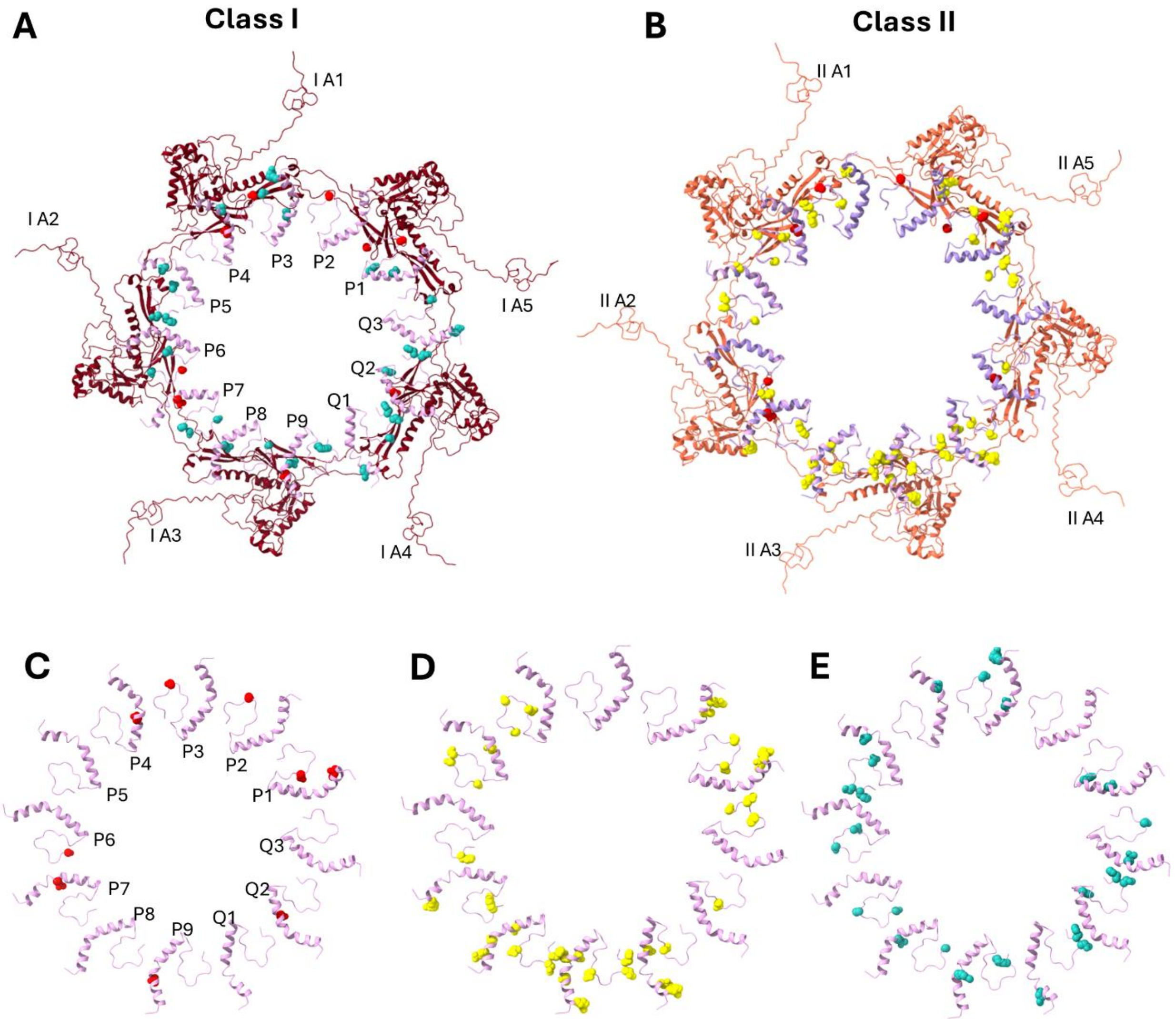

**Figure S5 - Residues on gp8 interacting with capsomers from 2 classes of tail assemblies.** A) Gp8 residues interacting with Chains A1-A5 from Class I assembly. B) Gp8 residues interacting with Chains A1-A5 from Class II assembly. C) N term residues colored red on gp8 interacting with chains A1-A5 from both Class I and II assemblies. D) Residues on gp8 colored red interacting only with Class I A1-A5 assemblies. E) Residues on gp8 colored light green interacting only with Class II A1-A5 assemblies.

**Table S1 – Cryo-EM data collection parameters**

| | |
|---|---|
| Magnification | 81,000 |
| Voltage (kV) | 300 |
| Microscope | TFS Titan Krios |
| Camera | K3 (counted) |
| Electron Dose | 45 $e^-$/ $Å^2$ |
| Defocus | -0.75 to -1.5 µm |
| Pixel size | 1.1 Å |
| Final pixel size | 1.5 Å (F-cropped) |
| Total movies collected | 32,103 |
| Movies used | 29368 |
| Total initial particles extracted | 1,037,050 |
| Total final particles used | 663,668 |

**Table S2 Cryo-EM map statistics**

| Details | | Particles | Symmetry | Resolution (Å) 0.143 FSC | EMDB codes |
|---|---|---|---|---|---|
| Class I | Capsid | 111,925 | I relaxed | 3.03 | EMD-78014 |
| | Tail | 100,440 | C1 | 3.03 | |
| Class II | Capsid | 113,545 | I relaxed | 3.03 | EMD-78027 |
| | Tail | 98505 | C1 | 3.03 | |
| Class III | Capsid | 113,982 | I relaxed | 3.04 | |
| | Tail | | | | |
| Class IV | Capsid | 113,493 | I relaxed | 3.02 | |
| | Tail | | | | |
| Class V | Capsid | 114,599 | I relaxed | 3.03 | |
| | Tail | | | | |

**Table S3 - Refinement and validation statistics for portal-tail complex from Class I**

| Assembly | Adaptor | Portal | Fiber trimer | Tail/Nozzle | Capsid |
|---|---|---|---|---|---|
| **PDB ID** | 37AN | | | | 37AT |
| **Composition** | | | | | |
| Chains | 12 | 12 | 3 | 6 | 6 |
| Atoms | 19680 (Hydrogens: 0) | 46644 (Hydrogens: 0) | 2722 (Hydrogen: 0) | 45498 (Hydrogen: 0) | 16473 (Hydrogen: 0) |
| Residues | Protein: 2248, Nucleotide:0 | Protein: 6036, Nucleotide:0 | Protein: 350, Nucleotide:0 | Protein: 5856, Nucleotide:0 | Protein: 2223, Nucleotide:0 |
| Water | 0 | 0 | 0 | 0 | 0 |
| Ligands | 0 | 0 | 0 | 0 | 0 |
| **Bonds (RMSD)** | | | | | |
| Length (Å) (# > 4s) | 0.003 (0) | 0.03 | 0.004 | 0.004 | 0.004 |
| Angles (°) (# > 4s) | 0.644 (0) | 0.702 | 0.9 | 0.579 | 0.613 |
| Molprobity score | 1.82 | 1.98 | 2.55 | 1.96 | 2.05 |
| Clashscore | 4.92 | 8.17 | 17.08 | 5.5 | 5.81 |
| **Ramachandran plot (%)** | | | | | |
| Outliers | 0.21 | 0.08 | 0 | 0.27 | 0.13 |
| Allowed | 3.55 | 4.04 | 4.94 | 5.75 | 4.07 |
| Favored | 96.25 | 95.87 | 95.06 | 93.98 | 95.75 |
| Rotamer outliers (%) | 2.53 | 2.17 | 4.26 | 2.22 | 3.77 |
| Cß outliers (%) | 0 | 0 | 0 | 0 | 0 |
| **Model vs. Data** | | | | | |
| CC (volume) | 0.79 | 0.8 | 0.74 | 0.88 | 0.85 |

**Table S4 - Refinement and validation statistics for Portal-tail complex from Class II**

| Assembly | Adaptor | Portal | Fiber trimer | Tail/Nozzle | Capsid |
|---|---|---|---|---|---|
| **PDB ID** | 37CZ | | | | 37AB |
| **Composition** | | | | | |
| Chains | 12 | 12 | 3 | 6 | 6 |
| Atoms | 19680 (Hydrogens: 0) | 46644 (Hydrogens: 0) | 2722 (Hydrogen: 0) | 45498 (Hydrogen: 0) | 16657 (Hydrogen: 0) |
| Residues | Protein: 2248, Nucleotide: 0 | Protein: 6036, Nucleotide: 0 | Protein: 350, Nucleotide: 0 | Protein: 5856, Nucleotide: 0 | Protein: 2250, Nucleotide: 0 |
| Water | 0 | 0 | 0 | 0 | 0 |
| Ligands | 0 | 0 | 0 | 0 | 0 |
| **Bonds (RMSD)** | | | | | |
| Length (Å) (# > 4s) | 0.003 (0) | 0.03 | 0.004 | 0.007 | 0.005 |
| Angles (°) (# > 4s) | 0.611 (0) | 0.786 | 0.9 | 0.683 | 0.684 |
| Molprobity score | 1.65 | 1.75 | 2.55 | 1.94 | 2.09 |
| Clashscore | 4.8 | 6.3 | 17.08 | 5.43 | 5.86 |
| **Ramachandran plot (%)** | | | | | |
| Outliers | 0.08 | 0.17 | 0 | 0.27 | 0.13 |
| Allowed | 2.68 | 3.89 | 4.94 | 6.33 | 4.56 |
| Favored | 97.24 | 95.94 | 95.06 | 93.39 | 95.31 |
| Rotamer outliers (%) | 1.8 | 1.45 | 4.26 | 1.9 | 3.79 |
| Cß outliers (%) | 0 | 0 | 0 | 0 | 0 |
| **Model vs. Data** | | | | | |
| CC (volume) | 0.85 | 0.77 | 0.74 | 0.88 | 0.85 |

**Table S5 Comparison of residue wise gp8:gp10 interactions in models from Class I and II assemblies based on PISA analysis.** (* indicates salt bridge. Highlighted residues indicate common gp8 residues which interact with different gp10 residues in Class I and II)

| **Class I** | |
|---|---|
| **gp8** | **gp10** |
| P1:Asp19 | a5:Gln329 |
| P1:Glu26 | a5:Lys58* |
| P1:Ser38 | a5:Arg49 |
| P1:Ser38 | a5:Asp50 |
| P1:Ser45 | a5:Thr56 |
| P2:Ser39 | b5:Ser126 |
| P3:Arg4 | a1:Glu120** |
| P3:Asp19 | a1:Gly335 |
| P3:Ser38 | a1:Val330 |
| P3:Ser39 | a1:Val330 |
| P3:Ser39 | a1:Pro328 |
| P4:Asp19 | a1:Thr56 |
| P4:Thr12 | a1:Asp50 |
| P4:Ser266 | b1:Tyr27 |
| P4:Gly262 | b1:Tyr27 |
| P5:Thr11 | b1:Gln329 |
| P5:Asp35 | b1:Asn334 |
| P5:Asn42 | a2:Val330 |
| P5:His43 | a2:Gln331 |
| P6:Asp35 | a2:Thr56 |
| P6:Ser39 | a2:Arg55 |

| **Class II** | |
|---|---|
| **gp8** | **gp10** |
| P1:4Arg | a5:Glu120* |
| P1:8Asn | a5:Glu120 |
| P1:Asp35 | a5:Gln331 |
| P1:Asp35 | a5:Asp334 |
| P1:Ser38 | a5:Val330 |
| P1:Ser39 | a5:Val330 |
| P1:Ser45 | a5:Ala325 |
| P2:Thr12 | a5:Lys54 |
| P2:Gln9 | a5:Asp50 |
| P2:Ser39 | b5:Lys40 |
| P3:Ser39 | a1:Asn344 |
| P3:Ser39 | a1:Asp324 |
| P3:Asn42 | a1:Gln331 |
| P3:His43 | a1:Gln331 |
| P3:Asn8 | b5:Gln329 |
| P3:Asp35 | b5:Asn334 |
| P4:Asp19 | a1:Lys58* |
| P4:Asp35 | a1:Lys58* |
| P4:Ser39 | a1:Arg55 |
| P4:Ser45 | b1:Ser35 |
| P5:Thr12 | b1:Lys127 |
| P5:Ser38 | b1:Val330 |
| P5:Ser38 | b1:Gln329 |
| P5:Ser38 | b1:Pro328 |
| P5:Ser39 | b1:Gln331 |
| P5:Ser39 | b1:Val330 |
| P5:Ser45 | b1:Asn334 |
| P6:His43 | a2:Thr56 |
| P6:Ser45 | a2:Lys58 |

| | |
|---|---|
| P6:Ser45 | b2:Phe36 |
| P7:Gln15 | b2:Glu120 |
| P7:Ser38 | b2:Val330 |
| P7:Ser39 | b2:Val330 |
| P7:Ser39 | b2:Gln331 |
| P7:Ser45 | b2:Asn334 |
| P8:Gln15 | a3:Gln331 |
| P8:Ser45 | a3:Thr56 |
| P8:Ser45 | a3:Lys58 |
| P9:Thr12 | a3:Lys317 |
| P9:Thr12 | a3:Arg55 |
| P9:Gln15 | b3:Glu37 |
| P9:Lys44 | b3:Glu120* |
| P10:Asn42 | b3:Pro328 |
| P10:His43 | b3:Gln329 |
| P10:Gln9 | b3:Gln329 |
| Q1:Ser39 | a4:Val332 |
| P11:Asp19 | a4:Thr56 |
| P11:Glu26 | a4:Lys58* |
| P11:Asn42 | b4:Glu120 |
| P11:His43 | b4:Glu120 |
| P12:Glu5 | b4:Arg49* |
| 12:Ser39 | b4:Gln329 |
| P12:Ser39 | a5:Asp111 |

| | |
|---|---|
| P6:Ser45 | a2:Thr56 |
| P7:Arg4 | a2:Gln204 |
| P7:Gln15 | b2:Glu37 |
| P7:Lys44 | b2:Glu120 |
| P8:Gln9 | a3:Gln329 |
| P8:Thr12 | a3:Gln329 |
| P8:Asp19 | a3:Asn334 |
| P8:Asp36 | a3:Asn334 |
| P8:Ser39 | a3:Val332 |
| P8:Asn42 | a3:Gln329 |
| P8:His43 | a3:Gln329 |
| P9:Arg4 | a3:Tyr252 |
| P9:Thr12 | a3:Arg49 |
| P9:Thr12 | a3:Gln329 |
| P9:Met16 | a3:Gly327 |
| P9:Asp19 | a3:Lys58* |
| P3:Ser45 | b3:His118 |
| P3:Asn42 | b3:Glu120 |
| P10:Arg4 | b3:Glu324** |
| P10:Asn8 | b3:Pro328 |
| P10:Asp19 | b3:Arg122** |
| P10:Asp35 | b3:Asn334 |
| P10:Ser38 | b3:Gln329 |
| P11:Asp19 | a4:Lys58 |
| P11:Asp19 | a4:Gln331 |
| P11:Asp19 | a4:Gln329 |
| P11:Asp35 | a4:Lys58 |
| P12:Arg40 | b4:Gln329 |
| P12:Asn42 | b4:Gln329 |
| P12:Ser45 | b4:Asn334 |

**Table S6 – Comparison of chain wise gp8:gp10 interaction in models from Class I and II**

| Class I | Class II |
|---|---|
| P1-a5 | P1-a5 |
| P2-b5 | P2-a5,b5 |
| P3-a1 | P3-b5,a1 |
| P4-a1,b1 | P4-a1,b1 |
| P5-b1,a2 | P5-b1 |
| P6-a2,b2 | P6-a2 |
| P7-b2 | P7-a2,b2 |
| P8-a3 | P8-a3 |
| P9-a3,b3 | P9-a3,b3 |
| P10-b3,a4 | P10-b3 |
| P11-a4,b4 | P11-a4 |
| P12-a5,b4 | P12-b4 |
